# Neuronal mechanisms of adaptive value coding in the amygdala

**DOI:** 10.1101/2024.10.13.618086

**Authors:** Julian Hinz, Mathias Mahn, Sigrid Müller, András Szőnyi, Tobias Eichlisberger, Andreas Lüthi

## Abstract

To address their needs, animals must optimize behavior by integrating external cues with internal state signals, such as hunger or thirst, environmental conditions, and past experiences. Accordingly, the perceived value of a reward varies depending on an animal’s current needs. Updating the specific value of rewards predicted by external cues depends on the basolateral amygdala (BLA). However, neurophysiological investigations have centered around the assumption of stimulus-invariant value representations and have not been able to account for behavioral findings that support the BLA’s ability to represent stimulus-specific value of distinct appetitive or aversive outcomes. To address how the BLA encodes and updates stimulus-specific value representations, we exposed head-fixed mice to distinct sets of gustatory rewards while tracking perceived value using lick microstructural analysis. Two-photon calcium imaging of BLA principal neurons revealed that the magnitude of the BLA population response scaled with perceived reward value and that the values of different rewards were encoded by distinct neuronal subpopulations. Moreover, reward representations rapidly re-scaled when mice were exposed to a reward that was larger than all previous rewards. Finally, value representations depended on an animal’s internal state as thirst selectively increased the responses to water rewards whereas aversive experiences strongly attenuated responses to sucrose. Our findings demonstrate that value representations in the BLA are stimulus-specific and highly adaptive to account for changes in relative reward value and to reflect an animal’s current affective and homeostatic state. This mechanism enables sensory-specific value updates necessary for state-adapted decision making and learning.

## Introduction

A major challenge for animals is to rapidly adjust their behavior to accommodate variable environmental conditions and need states. One of the main mechanisms to structure this interaction is associative learning in which actions or neutral sensory cues become predictive of appetitive or aversive outcomes. During associative learning, the value of the expected outcome is defined by its valence, which can be appetitive or aversive, and by its subjective importance or value.

One of the key brain areas for the acquisition, retrieval, and updating of associations of cues or actions with the current value of specific outcomes is the basolateral amygdala (BLA)^1,2^. Impairing BLA function prevents the formation and retrieval of both appetitive and aversive stimulus-specific associations^3–5^. Moreover, the BLA was found to be necessary to adapt behavioral responses whenever the perceived value of the expected outcome changed^6–9^. However, the BLA is not implicated in general motivation^10^ and contrary to other brain regions, optogenetic activation doesn’t cause excessive food consumption or strong place avoidance^11–13^. These findings suggest that behavioral adaptations are driven by dynamic changes in the representation of specific outcomes in the BLA. However, previous neurophysiological investigations of outcome representations in the BLA have so far suggested, sensory-invariant value representations^14–16^, which would not allow for flexibly updating value associations in a stimulus-specific manner and can’t account for behavioral evidence of sensory-specific value updating and learning. As stimulus specificity and dynamic value updating of outcome representations can only be characterized by exposing animals to a diverse set of rewarding outcomes, we performed two-photon calcium imaging in head-fixed mice consuming a range of appetitive gustatory stimuli. By measuring lick bout size, an established read-out for perceived value in experimental psychology^17,18^, we were able to quantify the relative values of these stimuli as a function of the rewards available and an animal’s internal state.

Our study reconciles previous findings at the behavioral and the neurophysiological level by demonstrating that the BLA represents the value of distinct appetitive outcomes in largely separate neuronal populations. Furthermore, we extend the concept of stimulus-specific reward value by showing that these representations are dynamically updated according to reward context, homeostatic and affective states, while population-level responses could be integrated to achieve a stimulus-invariant representation.

### BLA reward responses are locked to consumption onset

To study BLA activity during value assignment to innately valuable stimuli, we developed a head-fixed task in which food-restricted mice were presented with sets of unsignaled liquid rewards, delivered in a random sequence at random intervals, while recording general motor activity (running wheel motion), arousal (pupil dilation, whisker pad motion), and licking behavior (Fig. 1a). To minimize predictability and potential interactions between successive stimuli, we chose long inter-trial intervals of on average 20 s (Extended Data Fig. 1a). Quantification of licking behavior upon presentation with water or 20% sucrose demonstrated that animals continued to lick the reward spout long after the reward had been consumed, which takes about 3 licks (Fig. 1b-e and Extended Data Fig. 1b). Sucrose rewards resulted in significantly more licks per reward compared to water (Fig. 1d-e). The number of licks per reward was defined as the number of uninterrupted licks (inter lick interval < 500 ms) after reward delivery, also referred to as lick bout size. The difference in lick bout size was predominantly driven by a change in lick duration rather than frequency (Fig. 1d), as described previously^19^. To assess whether reward history influenced subsequent reward consumptions, we compared pupil dilation in trials with no change in reward and in trials in which there was a shift from water to 20% sucrose. We found no significant difference within the range of mean reward size fluctuations and therefore considered reward responses in subsequent trials to be independent from each other (Extended Data Fig. 1c-e).

**Fig. 1.**
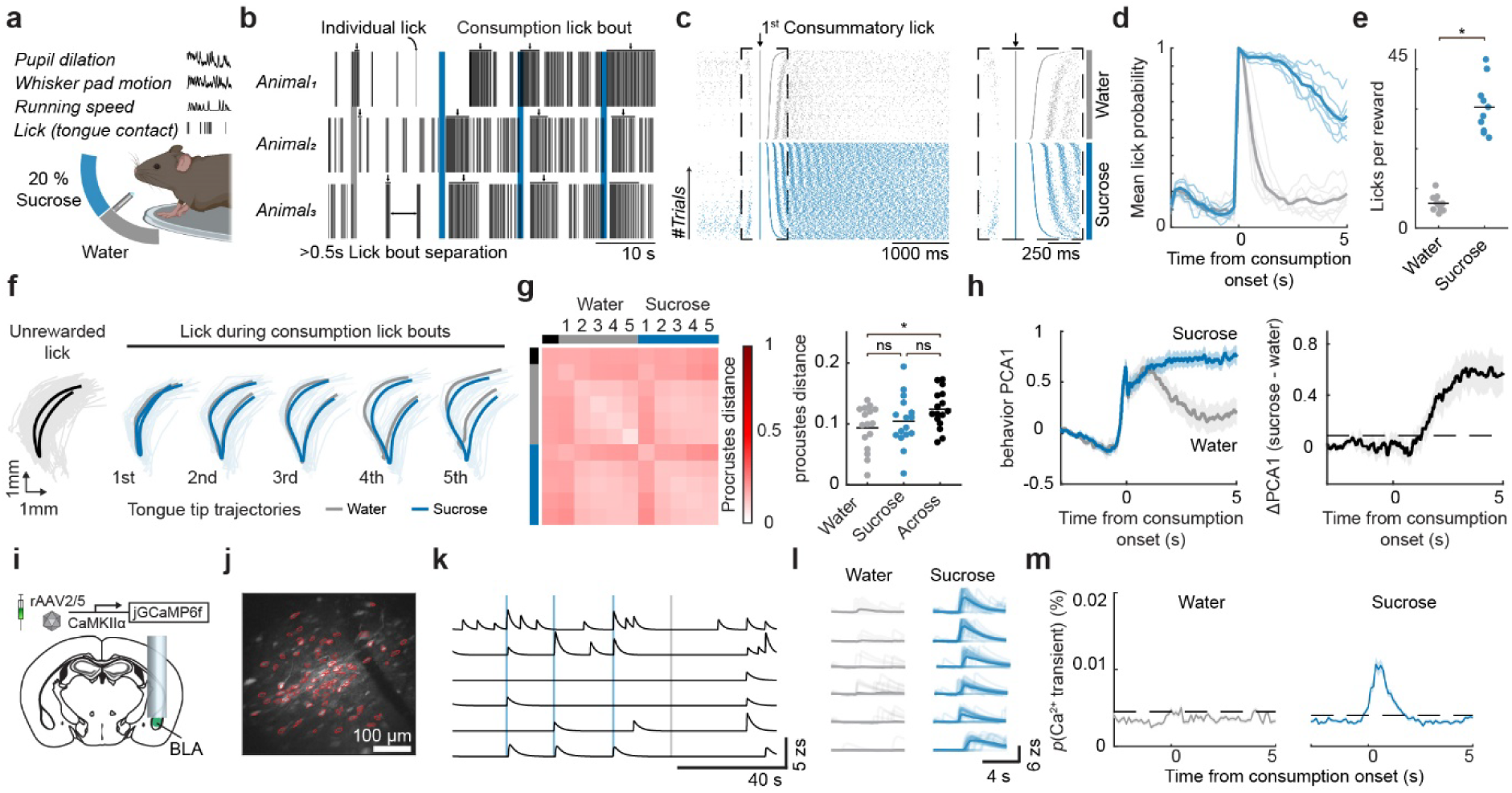
BLA activity is locked to reward consumption onset. **(a)** Schema of recorded variables during head-fixed behavioral paradigm with random reward deliveries. **(b)** Example licking from three animals consuming water and 20% sucrose rewards. **(c)** Lick pattern surrounding the first consummatory lick (*n* = 10 (animals) - sessions = 4). **(d)** Mean lick probability following consumption onset. **(e)** Mean number of uninterrupted licks (inter lick interval < 500 ms) following consumption onset for the different rewards (*p* = 0.0044, *n* = 10 (animals) - sessions = 4). **(f)** Example tongue tip trajectories during unrewarded licks as well as consummatory lick bouts whilst consuming water and sucrose 20%. Single interpolated lick trajectories with average trajectory overlaid. **(g)** Left: matrix comparing the procrustes distance of different lick categories (reward type and lick number within lick bout). Right: Quantification of interpolated lick trajectory shape differences using Procrustes distance within a lick category (same reward and lick within lick bout) compared to across the two reward types (*p* = 0.0014; Within water to within sucrose: *p* = 0.33, within water to across: *p* = 0.0011, within sucrose to across: *p* = 0.071; *n* = 16 (*sessions*) - *animals* = 4) **(h)** PCA1 of the combined behavioral read-outs of whisker pad motion, pupil dilation and running speed (left) and the delta between the combined read-outs between water and sucrose 20% (right). Dashed line indicates a 3 standard deviation increase from baseline. **(i)** Schema of the viral and surgical strategy for two-photon recordings in the BLA with an example recording plane shown in **(j).** Red outlines indicate the spatial footprints of the extracted neurons. **(k)** Example traces of 6 simultaneously recorded neurons. **(l)** Single (opaque) and mean (solid) reward-triggered activity of the neurons in **(k)**. **(m)** Calcium transient probability in response to reward consumption (*n* = 1846 (*neurons*) - *animals* = 10, *sessions* = 4). Dashed lines indicate a 3 standard deviation increase from baseline transient probability.

To address whether licking behavior is reward-specific, we analyzed tongue movements at higher temporal resolution using video tracking at 194 Hz (Supplementary Video 1). Tongue motion was stereotypical, with probing tongue movements to check for reward availability and a transition to a lapping motion after reward detection (Fig. 1f). Comparing lick trajectories revealed that licks in response to water were more similar to each other than licks in response to 20% sucrose (Fig. 1g), while 20% sucrose licks were as similar to each other as to water. To test whether such lick pattern differences could systematically bias our neuronal recordings, we quantified if differences in lick pattern scale with sucrose concentrations differences and found that they did not (Extended Data Fig. 1f-h). To further assess at which point consumption behavior started to differ, we compared the first principal component of the combined behavioral metrics (whisking pad motion, running speed, and pupil dilation) for water and sucrose consumption lick bouts (Fig 1h - left).

We found that the behavior started to significantly deviate 1.28 s after consumption bout onset (Fig. 1h - right), indicating that during this time window neural activity is likely related to the sensory content or value of the stimulus, rather than to differences in motor patterns^20^.

To characterize the neuronal representation of reward value, we expressed the calcium indicator GCaMP6f in principal neurons in the BLA (Fig. 1i and Extended Data Fig. 2) and imaged neurons through a gradient refractive index (GRIN) lens during the task using a two-photon microscope. Aligning the neuronal responses to the onset of reward consumption indicated that BLA principal neurons showed calcium responses time-locked to consumption onset (Fig. 1 i-l). Rather than spanning the entire time of a given lick bout, reward responses were primarily onset responses. The probability of calcium transient onset was elevated above baseline during the first 1.5 s after consumption onset (Fig. 1M). This timing makes it unlikely that the motor pattern differences described above contribute to the measured differences in neuronal activity. To further control for a possible contribution of lick-related motor signals, we sub-selected trials with similar lick bout sizes for both rewards and found that reward response differences were maintained (Extended Data Fig. 3a-c).

### BLA population activity reflects perceived reward value

To address how the BLA represents the value of rewards across a range of gustatory stimuli, we exposed mice to three different reward sets (Fig. 2a, f, k). Lick bout sizes were normalized to the average lick bout size in response to 10 µl 20 % sucrose reward (normalized licking), which was present in all reward sets. This analysis showed that normalized licking scaled with the animals’ preference for higher sucrose concentrations (Fig. 2b) and larger volumes at a given concentration (Fig. 2g). We also observed differential licking for distinct rewards in a set composed of rewards containing different nutrients (milk, flavored milk, and sucrose; Fig. 2l), where value assignment was unclear a *priori*. As previous work used lick bout size as a proxy for relative value assessment^21^, we conclude that licking behavior can also be used to measure the perceived value of unpredicted gustatory rewards in our head-fixed task.

**Fig. 2.**
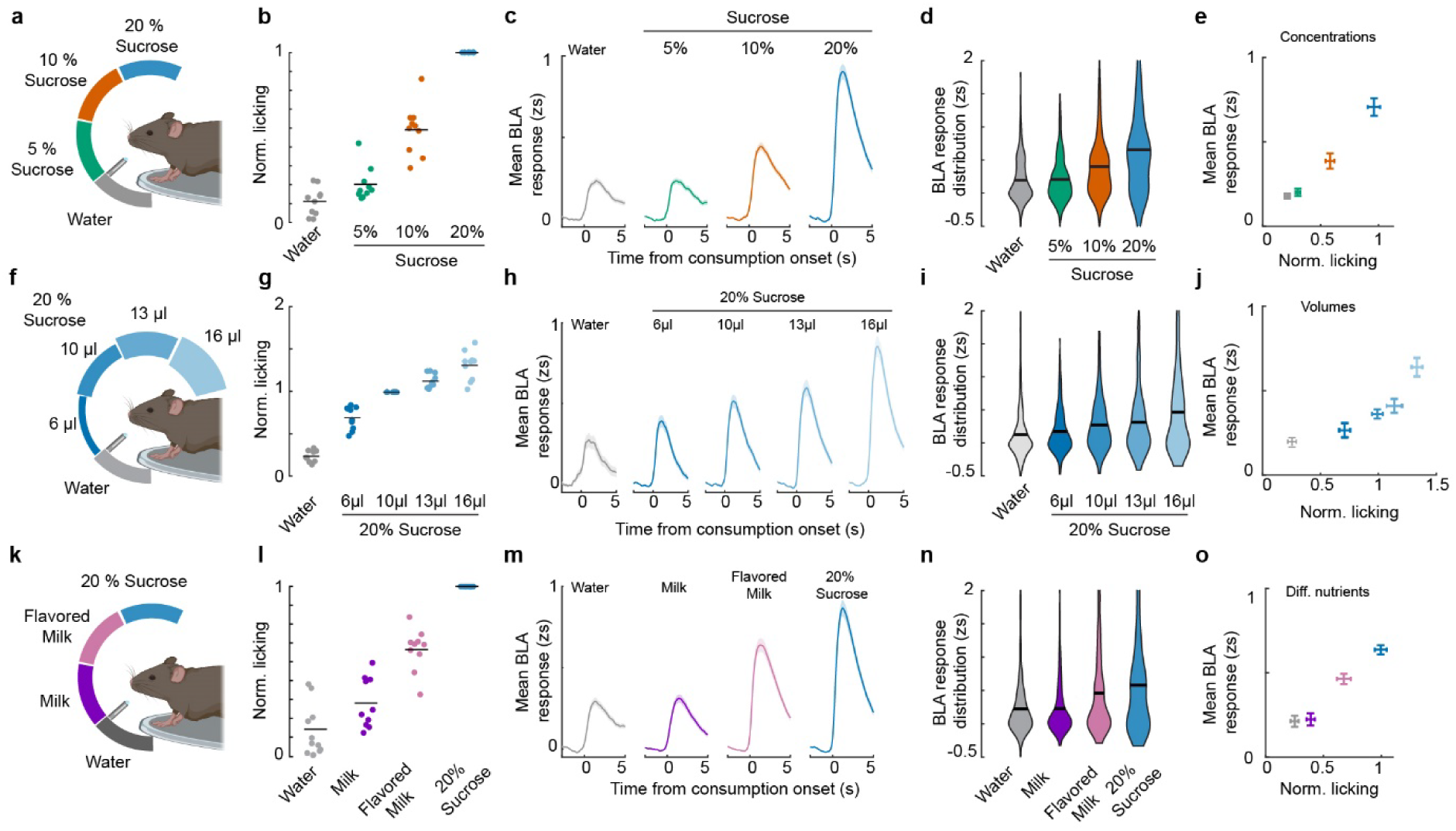
The BLA represents the value of gustatory rewards at the population level. **(a-e)** Characterization of reward concentration effects. **(a)** Schema of the randomly provided sucrose concentrations during the behavioral session (10 µl per reward). **(b)** Number of licks normalized to the 10 µl, 20% sucrose response in this session (*p* = 9.29*10^(−15), *n* = 10 (*animals*), *sessions* = 4). **(c, d)** Time-resolved mean population response of active BLA neurons **(c)** with corresponding quantification (*p* = 1.81*10^(−27), *n* = 427 (*neurons*); *animals* = 10, *sessions* = 4) **(d)**. **(e)** Average mean BLA response plotted against the mean normalized licking for the Volume session. **(f-j)** Characterization of reward volume effects. **(f)** Schema of the randomly provided volumes of 20% sucrose during the behavioral session. **(g)** Number of licks normalized to the 10 µl, 20% sucrose response in this session (p = 1.25*10^(−21), *n* = 10 (*animals*) - *sessions* = 2). **(h, i)** Time-resolved mean population response of active BLA neurons **(h)** with corresponding quantification (*p* = 0.013, *n* = 185 (*neurons*) - *animals* = 10, *sessions* = 2) **(i)**. **(j)** Average mean BLA response plotted against the mean normalized licking for the session. **(k-o)** Characterization of reward type effects. **(k)** Schema of the randomly provided reward types during the behavioral session (10 µl per reward). **(l)** Number of licks normalized to the 10 µl, 20% sucrose response in this session (*p* = 6.86*10^(−10), *n* = 10 (*animals*) - *sessions* = 3). **(m, n)** Time-resolved mean population response of active BLA neurons **(m)** with corresponding quantification (p = 5.62*10^(−19), *n* = 510 (*neurons*) - *animals* = 10, *sessions* = 4) **(n)**. **(j)** Average mean BLA response plotted against the mean normalized licking for the session.

For each reward set, we observed an increase in mean BLA activity across session averaged single neuron responses for preferred rewards (Fig. 2c-e, h-j, m-o). Due to the biophysical properties of the calcium sensor, full differentiation of the sources for this increased activity is not possible^22^. It is however likely that it reflects a combination of the recruitment of more neurons, an increase in single-neuron response amplitude and/or response probability (Extended Data Fig. 3e-g). A positive correlation of the mean neuronal response magnitude with normalized licking was also observed when subdividing responses based on lick bout size alone (Extended Data Fig. 3h) and when subdividing for individual rewards and lick bout size (Extended Data Fig. 3i). Thus, consistent with previous studies concluding that BLA reward representations are stimulus-invariant^15,16^, our results demonstrate that mean BLA activity correlates with lick bout size independent of the currently consumed reward, and that perceived reward values are represented in a stimulus-invariant manner at the population level.

### Neuronal subpopulations encode stimulus-specific reward value

The monotonically increasing relationship of perceived value and population response could be a result of stimulus-invariant value neurons that each have a monotonically increasing relationship with reward value or arise through the combination of subpopulations of neurons specifically responding to distinct rewards.

To distinguish between these two possibilities, we performed a regressor analysis to test whether neurons had stimulus-invariant responses by correlating mean responses with a regressor containing all stimuli and comparing it to the best correlating individual stimulus regressor (*see* methods, Fig. 3a). In the sessions in which mice were exposed to sucrose solutions of different concentrations and volumes and in which stimulus features were very similar, correlations for the best maximum individual regressor were slightly better than the complete stimulus regressor. This difference was increased during the session in which mice were exposed to distinct stimuli containing different nutrients, where we found a strong bias towards neurons correlating better with the maximum individual stimulus regressor (Fig. 3b, c). This finding indicates that individual neurons had stimulus-specific responses. To test this further, we compared the number of neurons that were significantly responding to only one reward and found that most neurons were responding exclusively to a single reward, with higher overlap during sessions with higher reward similarity (Fig. 3d-f). To understand whether the observed findings were only true for session averages or also observable at the individual trial level, we calculated population vector (PV) correlations (Fig. 3g). Comparing the across vs. within PV-stimulus correlations revealed that even at the single trial level the population response was significantly different for sessions in which different rewards were consumed (Fig. 3h).

**Fig. 3.**
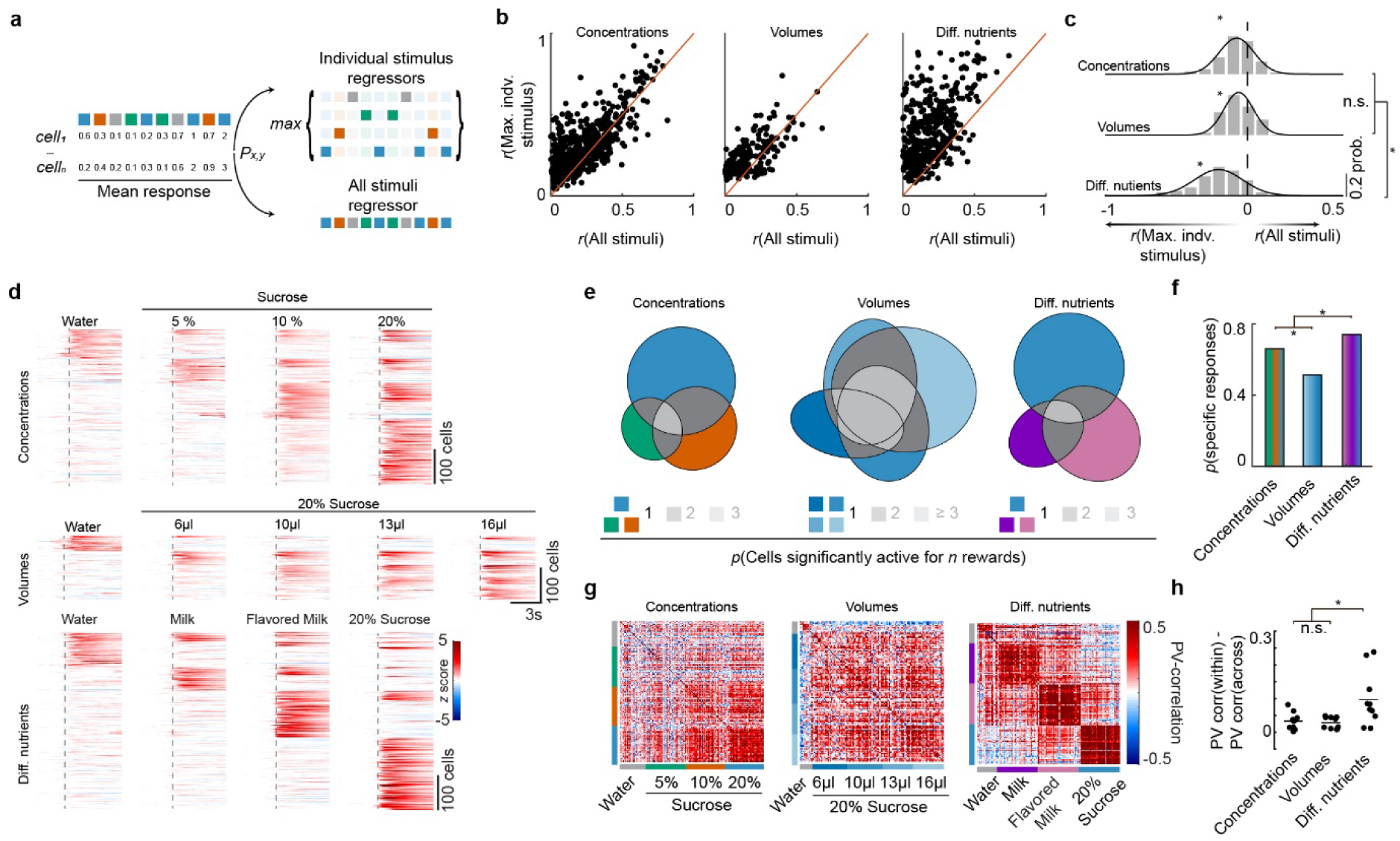
Distinct BLA sub-populations track stimulus-specific value. **(a)** Schema of linear regression analysis performed to characterize single neuron tuning. **(b)** Correlation of individual neuron responses with lick bout size and the respective stimuli in the Concentration (left), Volume (middle) and Diff. nutrient (right) sessions with corresponding quantification in **(c)** (Conc: *p* = 3.98*10^(−187), Vol: *p* = 6.13*10^(−81), Diff. nutrients: *p* = 5.58*10^(−223); Conc.-Vol. p > 0.5, Conc. - Diff. nutrients p = 5.45*10^(−39), Diff. nutrients - Vol. p = 2.72*10^(−23); - Concentrations: n = 427 (*neurons*) - *animals* = 10, *sessions* = 4; Volume: *n* = 185 (*neurons*); *animals* = 10, *sessions* = 2; Diff. nutrients: *n* = 510 (*neurons*) - *animals* = 10, *sessions* = 4). **(d)** Time-resolved activity of all significantly modulated neurons across the different behavioral sessions. **(e)** Venn diagrams displaying the reward specificity of individual neurons across different reward sessions with corresponding quantification of neurons active to only one reward shown in **(f)** (Conc. - Vol. p = 0.0018, Conc. - Diff. nutrients p = 0.0209, Diff. nutrients Vol. p = 3.83*10^(−8), Conc: n = 427 (*neurons*) - *animals* = 10, *sessions* = 4; Vol: *n* = 185 (*neurons*) - *animals* = 10, *sessions* = 2; Diff nutrients: *n* = 510 (*neurons*) - *animals* = 10, *sessions* = 4). Shaded regions depict SEM. **(g, h)** Trial-to-Trial correlation of the population vector (PV) for 3 example sessions of the types: Concentration, Volume, and Diff. nutrient with corresponding quantification of within reward vs. across reward PV-correlation for the individual sessions in **(h)** (Conc. - Vol. p = 0.97, Conc. – Diff. nutrients. p = 0.02, Diff. nutrients - Vol. p = 0.011, Conc.: *n* = 10 (*animals*) - *sessions* = 4; Vol.: *n* = 10 (*animals*) - *sessions* = 2; Diff. nutrients: *n* = 10 (*animals*) - *sessions* = 4).

Taken together the presented data demonstrate that the BLA maintains stimulus-specific value representations at the single neuron level, while the mean population response represents the relative reward value indiscriminate of stimulus identity.

### BLA neurons dynamically encode value

To address how BLA reward value representations change upon exposure to a preferred reward and to expand on previous work focusing on individual neurons^23^, we first performed a cross-session comparison of the neural response to the same stimulus (10 µl of 20% sucrose) in the presence and absence of a better, preferred reward. This comparison (Fig. 4a) revealed that animals licked less for the same reward when they experienced a larger 16 µl reward of the same concentration in the same session, while responses for water were not affected (session II; Fig. 4b). Similarly, we found that BLA responses were reduced for the 10 µl reward (session II) and that the 16 µl reward, at the time the most valuable available reward (session II), evoked a similar response as the 10 µl reward in sessions where mice were presented with 10 µl as the highest reward (session I; Fig. 4c, d). To test whether reward response re-scaling also occurs within the same experimental session, we performed an additional experiment where we alternated two different reward scales between blocks (Fig. 4e and Extended Data Fig. 2c), an experimental design known as simultaneous contrast^24^. Consistent with the previous findings, we observed larger lick bout sizes (Extended Data Fig. 6e) and a larger BLA population response to the same 20% sucrose reward within a reward block that did not contain 40% sucrose compared to blocks with 40% sucrose rewards available (Fig. 4f). Analyzing block transitions showed that BLA population responses adapted within a single session. 20% sucrose responses rapidly adapted within the blocks (5-10 min.) and recovered to baseline levels after termination of the 40% block (Fig. 4g, h). This effect was mediated by a group of neurons (Extended Data Fig. 4a) with stimulus-specific responses, consistent with our previous observations (Extended Data Fig. 4b). Within-session adaptation suggests an integration of local reward history, which we didn’t observe in previous experiments (Extended Data Fig. 1c-e). This discrepancy is likely explained by the higher mean reward fluctuations in the simultaneous contrast session (Extended Data Fig. 4c-d), an interpretation supported by control experiments in which we introduced an equal number of 20% sucrose trials instead of 40% sucrose (Extended Data Fig. 4f), where we did not observe an adaptation between the different blocks (Extended Data Fig. 4g-j).

**Fig. 4.**
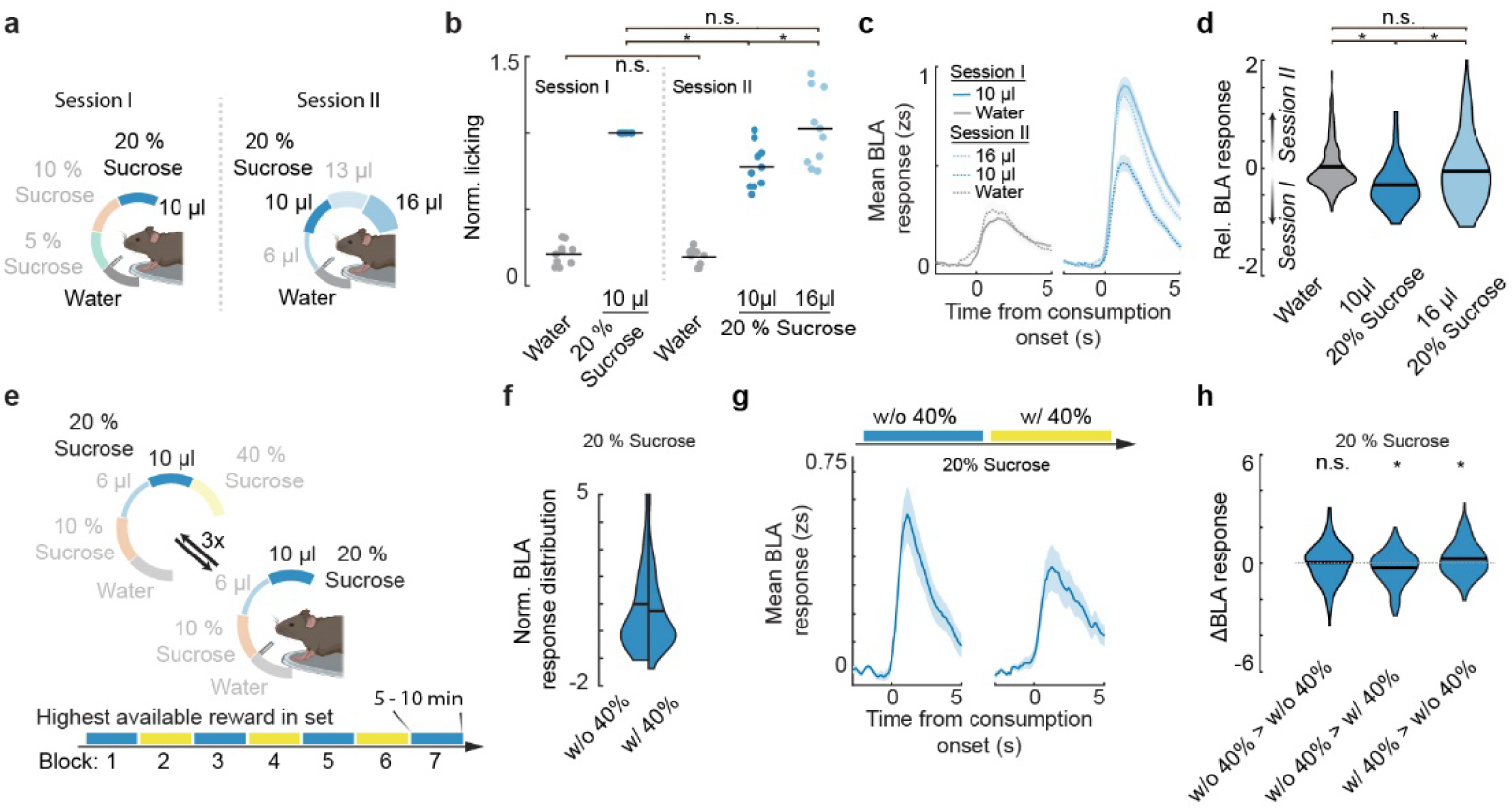
BLA maintains coherence of value representation by re-scaling upon exposure to a new reward environment. **(a)** Schema of the sessions, as presented in Fig. 2. **(b)** Number of licks normalized to the 10 µl 20% sucrose response in the concentration session (Session I) on a per animal basis (Conc. 10 µl - Vol. 10 µl: *p* = 0.0011; Vol. 16 µl - Vol. 10 µl: *p* = 0.0019; Conc. 10 µl - Vol. 16 µl: *p* > 0.5; Conc. water - Vol. water: *p >* 0.5; *n* = 10 (*animals*) - *session* = 4/2). **(c)** Time-resolved mean BLA response to rewards in the volume (dashed line) and concentration session (solid line) **(d)** with corresponding quantification of the relative BLA response of significantly responding neurons in the volume session (session II) subtracted from the mean response of significantly active neurons in the concentration session (session I) of the corresponding rewards depicted in **(c)** (water - 10 µl: p = 9.72*10^(−10); 10 – 16 µl: p = 2.33*10^(−6); water – 16 µl: *p* = 0.013; *n* = 185 (*neurons*) - *animals* = 10, *sessions* = 4/2). **(e)** Schema of the experimental design for simultaneous contrast characterization. Animals were given alternating blocks of rewards including (yellow line) or excluding (blue line) 40% sucrose. **(f)** Split violin plots without (left) and with (right) 40% sucrose present in the current block (*p* = 0.029, *n* = 104 (*neurons*) - *animals* = 4, *sessions* = 3). **(g)** Time-resolved mean BLA response to 20% in blocks without (blocks 1, 3, 5, 7) and the blocks with 40% sucrose (blocks 2, 4, 6). **(h)** Quantification of the BLA response change between 20% blocks (blocks 1-3, 3-5, 5-7, left), transitions from 20% blocks to 40% blocks (1-2, 3-4, 5-6, middle) and transitions from 40% blocks to 20% blocks (2-3, 4-5, 6-7, right) (p > 0.5; *p* = 0.01; *p* = 0.005; *n* = 218/174/176 (*ΔNeuron activity*) - *animals* = 4, *sessions* = 3, *blocks* = 3, *neurons* = 104). Error bars and shaded regions depict SEM.

### Value representations reflect homeostatic and affective states

To examine whether the BLA response scaling does not only take the distribution of all available rewards into account but also incorporates changes in an animal’s internal state, we first measured reward responses in acutely water-restricted animals. At the behavioral level, we observed a strong and specific increase in the lick response to water, indicating a shift in the relative value of water in comparison to sucrose (Fig. 5a-c). Accordingly, the BLA population response to water was specifically increased in water-restricted animals, resulting in an equal BLA population response magnitude to water and sucrose rewards (Fig. 5d-f). This increased water response was predominantly driven by an increase in the number of neurons showing a water-specific response that we did not observe in non-water-restricted animals (Fig. 5e, f).

**Fig. 5.**
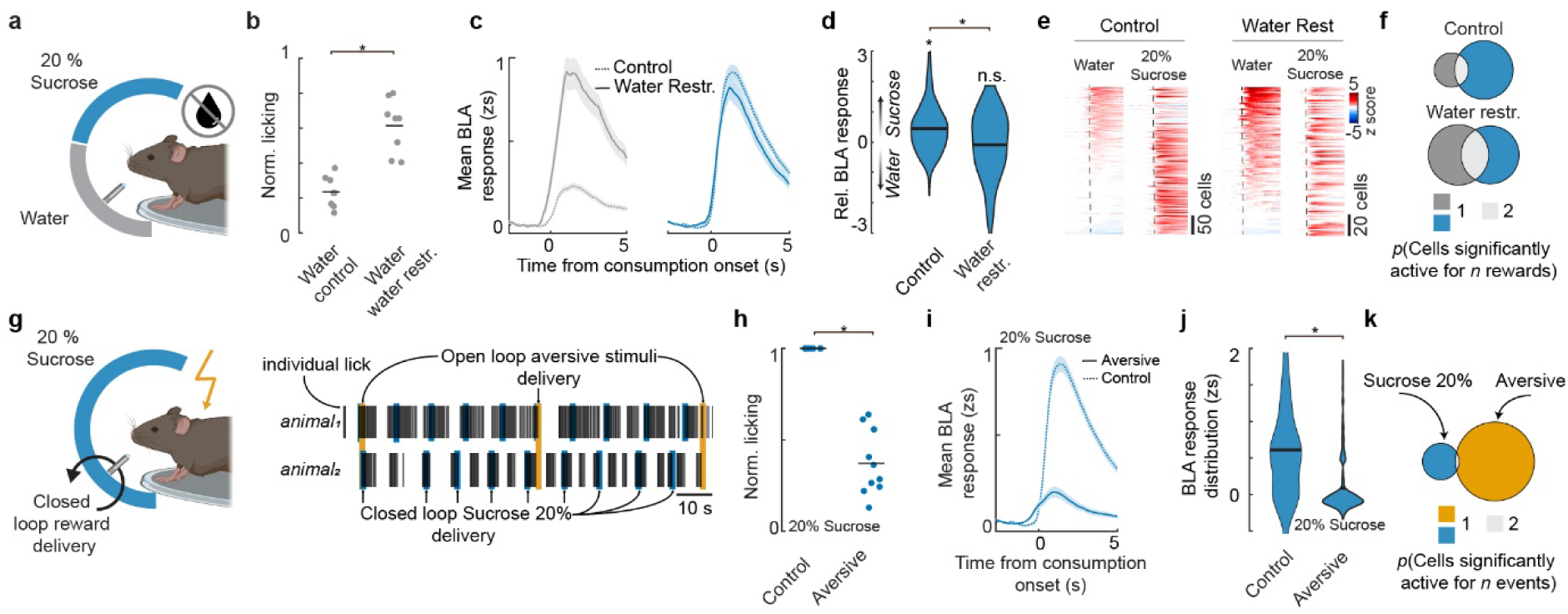
Value representations in BLA are internal state dependent. **(a)** Schema of water deprivation session design. **(b)** Number of licks normalized to the 10 µl, 20% sucrose response in the session exposing animals to different sucrose concentrations. Comparison of the licking response to water during the water deprivation session with the concentration sessions (*see* Fig. 2b; *p* = 0.00022, *n* = 10 (*animals*) - *sessions* = 4/1). **(c)** Time-resolved mean BLA response to different rewards of significantly active neurons sorted by mean activity to water for water restriction (left) and control condition (right). **(d)** Quantification of the difference of all significantly active neurons for water responses in the control condition (concentration session) and water restriction session (FR - WR: *p* = 2.4*10^(−10); FR: *p* = 8.48*10^(−36); WR: *p* = 0.27; FR: *n* = 427 (*neurons*); *animals* = 10, *sessions* = 4, WR: *n* = 104 (*neurons*); *animals* = 10, *sessions* = 1). **(e)** Time resolved activity of all BLA neurons during water restriction, and control session (*see* Fig. 2b). **(f)** Visualization of overlap of significantly water and 20% sucrose responsive neurons in control session (upper panel) and water restricted session (lower panel). Normalized to the fraction of significantly 20% sucrose active neurons in the control session. **(g)** Schema of aversive session design. **(h)** Licking response comparing the licking during control session with this session (*p* = 4.14 *10^(−36); *n* = 10 (*animals*) - *sessions* = 3). **(i, j)** Time-resolved mean BLA response to 20% sucrose reward during the aversive session (solid line) and the food restricted session (dashed line; see Fig. 2b) with corresponding quantifications (*p* = 2.33*10^(−31), *n* = 427/258 (*neurons*) *animals* = 10, *sessions* = 4/3) **(j)**. **(k)** Overlap of significantly responding neurons responding to aversive events and 20% sucrose. Shaded regions depict SEM.

Finally, we examined how a negative affective state would influence reward value representations by exposing mice to unpredicted aversive stimuli randomly interspersed throughout the session (random ITI 45 – 60s, Fig. 5h). We further modified the paradigm to trigger reward delivery upon spout licking, allowing us to dissociate motivational drive from assigned value^25^, which could both be influenced by repeated aversive exposures. Aversive experiences led to a marked decrease in the lick bout size after consumption but did not result in a strong reduction of reward seeking behavior, as quantified by lick-triggered reward deliveries (Fig. 5h-j and Extended Data Fig. 5a-d). While reward-seeking behavior was only partially affected and the BLA strongly responded to aversive stimuli (Extended Data Fig. 5e, f), we observed a strong attenuation of neuronal sucrose responses despite continued consumption (Fig. 5k, l). Reward-responsive neurons were not overlapping with aversive-responsive neurons in this experiment, suggesting that the investigated appetitive value representation is likely valence specific (Fig. 5m).

## Discussion

Here we show that the BLA represents the perceived value of stimuli and that this effect is not a mere reflection of general arousal or systematic differences in licking microstructure. Furthermore, we find that this stimulus-invariant value response emerges from largely separate neuronal subpopulations specifically activated by distinct gustatory stimuli. Exposing mice to larger rewards within the same stimulus set resulted in re-scaling of reward representations, demonstrating that value representations are relative and maintained through rapid adaptation of neural response magnitudes. Finally, we show that changes in the internal state can bi-directionally modulate the magnitude of value responses in the BLA. These data thus support a pivotal role for the BLA at the interface between stimulus-specific sensory processing and stimulus-invariant value computation.

Our finding of consumption onset locked responses is in agreement with previous reports of BLA activity in response to reward exposure^26^. However, we did not observe a reduction of consumption-onset locked responses across time^14^, potentially due to the lack of predictability of reward availability in our paradigm. Interestingly and contrary to reports in other brain areas, we find that single neurons in the BLA show stimulus-specific responses and are not dominated by orofacial movement (Extended Data Fig. 3, ^20^).

Previous studies have reported the existence of a stimulus-invariant value code in BLA^15,16,26^ which we find only at the population level. We extend these findings by demonstrating that even though the total population does indeed encode stimulus-invariant reward value, this representation arises from stimulus-specific value representations that vary in number of neurons, magnitude and response probability according to the perceived value of a given specific stimulus. The high stimulus specificity even for very similar rewards like different sucrose concentrations cannot be sustained with increasing stimulus set size and would lead to suboptimal coding^27^. Due to the overlap of neurons coding for sugar and fat-based rewards (Fig. 3) it is likely that, at least in the gustatory domain, the BLA projects stimulus features onto relevant axes, such as carbohydrate, protein, fat, and mineral content.

Our discovery of stimulus-specific response properties of BLA neurons allows to resolve a longstanding disconnect between behavioral studies, showing the necessity of BLA function when the value of specific rewards needs to be updated, for example during outcome devaluation, revaluation, reinstatement, conditioned taste aversion or sensory specific associations^6–10^ and neurophysiological reports of stimulus-invariant valence responses in the BLA^15,16,26^. Our findings are well aligned with recent work from our lab, describing the transfer of specific stimulus value to instrumental actions and the necessity for this representation for actions execution^28^ and are supported by holographic stimulation of BLA neurons responsive to sucrose, leading to an increase in the perceived value of sucrose^29^.

Our findings are also complementary to the characterization of learned responses to predicting neutral stimuli (such as tones) that are paired with appetitive or aversive stimuli^30^ and suggest a potential mechanism for the rapid update of sensory-specific associations ^3–5^. The adaptation of single neurons to the best currently available stimulus in the BLA have been reported in primates^23,31^ using different reward volumes. Here we show that this adaptation is more general and applies to volumes as well as concentrations. Moreover, we demonstrate that neurons adapt within minutes and recover the original representation once the previous reward set is presented.

Thirst modulated neuronal activity has been reported throughout the brain^32^ and it is yet unclear how the subfornical organ drives changes in value assignment. Given our results that show that water responses in the BLA are strongly increased upon water deprivation, and in the light of previous reports on the specific role of the BLA in updating stimulus-specific action and Pavlovian associations^8,10^, we speculate that the BLA is a key site in implementing this homeostatic-state dependent update.

The impact of an aversive state on food consumption has been observed at the level of the hypothalamus^33^, directly impacting the activity of hormonal driven hunger signals of AgRP neurons. Due to the unchanged high motivation of animals exposed to aversive stimuli to ingest food (Extended Data Fig. 5), it is unlikely that the decrease in activity in BLA following reward consumption is a consequence of the altered hunger signal broadcasted to the brain though the above-mentioned hypothalamic system, and strongly indicates that BLA reward representations are strongly modulated by an animal’s affective state.

Our results complement findings in lateral hypothalamus (LHA), a major target of BLA projections, that demonstrated that stimulus unspecific appetitive behaviors can be driven with optogenetic stimulation^34^, thus suggesting that the BLA is situated at the interface between stimulus-specific sensory processing and stimulus-invariant motivation. These findings also align well with observations in humans, where the amygdala has been found to be important for the integration of value, while its dysregulation has been linked to affective disorders accompanied by anhedonia^35–37^, suggesting an important role for the BLA in mediating value assignment under physiological conditions.

## Acknowledgments

The authors thank Yue Zhang, Martin Zeller, Nikolaos Karalis, Julien Courtin and all members of the Lüthi laboratory for discussion about project design and feedback on earlier versions of the Manuscript. We thank P. Argast and P. Buchmann for technical assistance, G. Ferrand, B. Heller-Stilb and the animal caretakers for help with animal husbandry and Charlotte Sonseson (Bioinformatics, FMI) for help with the statistical analysis. This work was supported by the Novartis Research Foundation, the European Research Council (ERC) under the European Union’s Horizon 2020 Research and Innovation Program (Grant Agreements 669582 to A.L. and under the Marie Skłodowska-Curie grant agreement 844492 to M.M.), the Swiss National Science Foundation (310030B_170268; TMAG- 3_209270; all to A.L. and Ambizione grant agreement No PZ00P3_209032 to M.M.), and the European Molecular Biology Organization (EMBO-ALTF-233-2020 to A.S.).

## Authors contributions

Conceptualization: JH, MM, AL

Methodology: JH, MM, AS

Investigation: JH, MM, SM, AS, AL

Visualization: JH, MM

Funding acquisition: MM, AS, AL

Project administration: JH, AL

Supervision: AL

Writing – original draft: JH, MM, AL

Writing – review & editing: JH, MM, AL

All authors have read and agreed to the published version of the manuscript.

---

**Extended Data Fig. 1:**
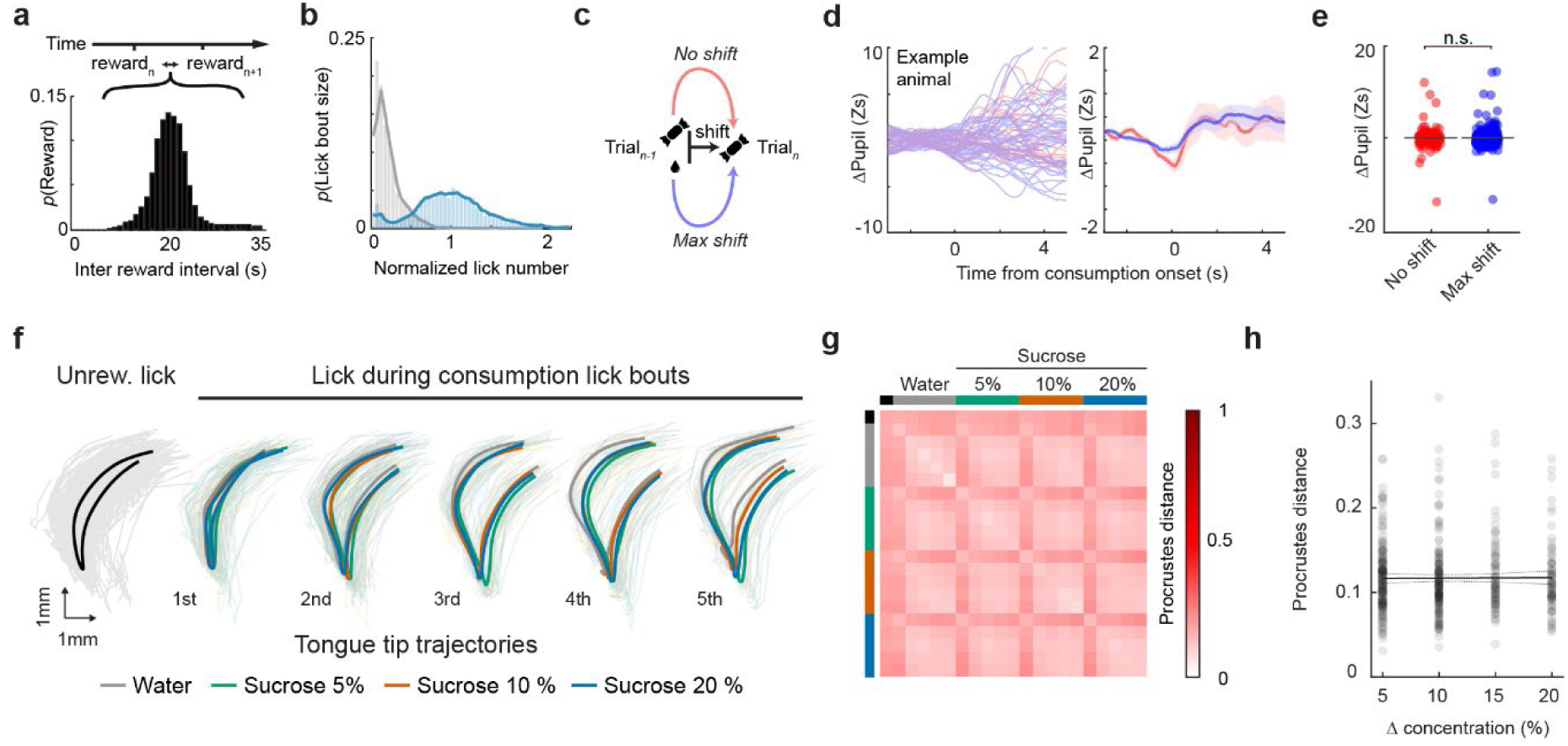
Animals show no measurable signs of reward history effect or concentration dependent lick pattern difference. (a) Distribution of inter-reward-intervals between reward deliveries during the task. (b) Distribution of lick number, normalized to the licking of each animal to 20% Sucrose (*n* = 5285 (consummatory lick bouts) - *animals* = 10, *sessions* = 4). (c) Schema of the three evaluated cases (d) Pupil trajectories of an example animal of 20% sucrose consumption with preceding water or 20% sucrose trial (left) and average across animals of the three conditions (right), with the corresponding quantification in (p =; n =) (e). (f-h) Same analysis as in Fig. 1 f extended to different sucrose concentrations to determine whether lick trajectory differences scale with reward value difference. (f) Example tongue tip trajectories during unrewarded (check licks) as well as consummatory lick bouts in response to reward types. Single lick trajectories with average trajectory overlaid. (g) Quantification of interpolated lick trajectory shape differences using Procrustes distance within a lick category (same reward and lick within lick bout) compared to other categories. (h) There is no dependence of lick trajectory dissimilarity and reward concentration difference (slope: 6*10^(−5) ± 3.8*10^(−4); *p*: 0.87).

**Extended Data Fig. 2:**
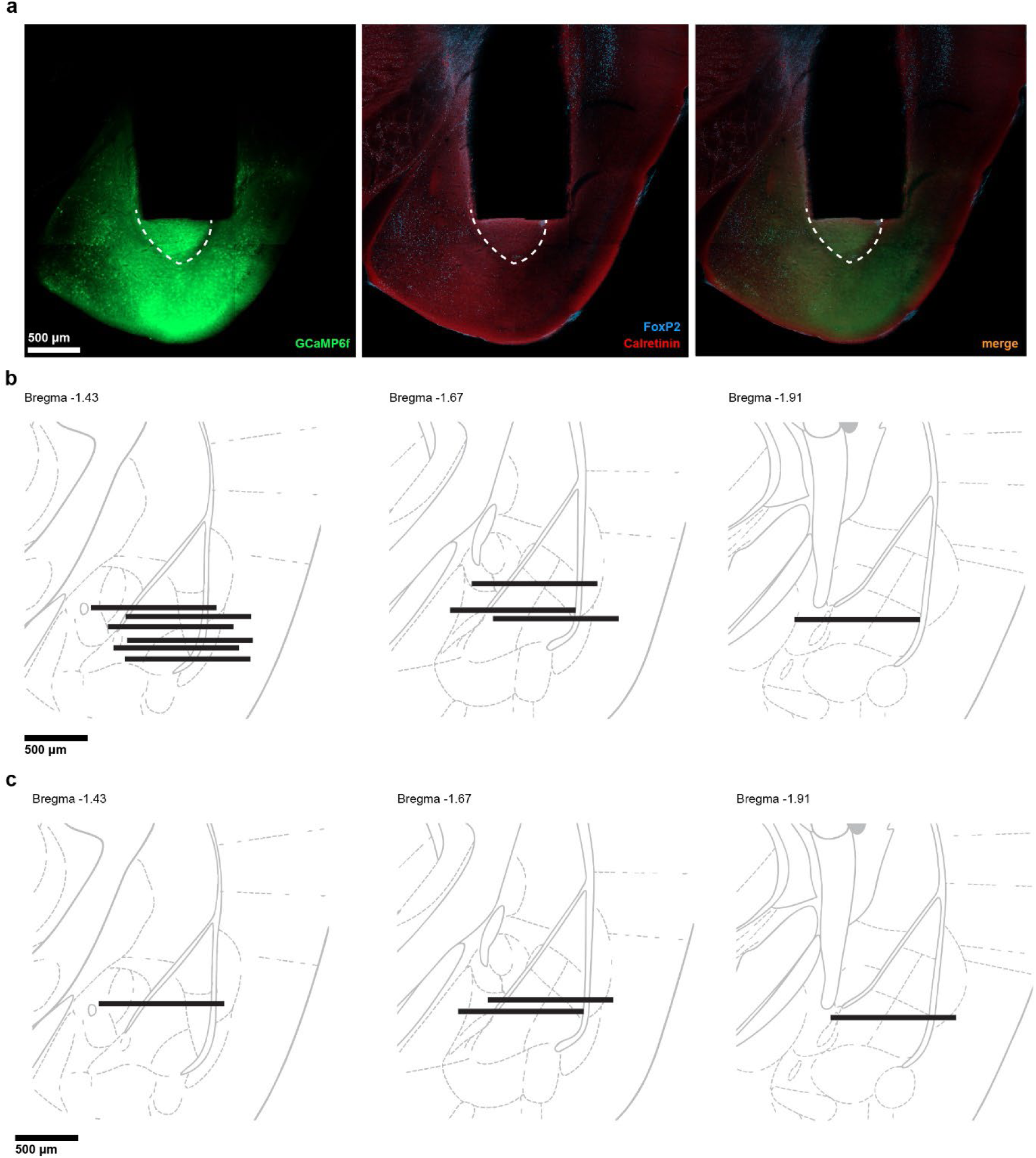
GRIN lens placements. To allow for reliable brain registration despite the GRIN lens induced deformations, we performed stainings against FoxP2 and Calretinin as landmarks. **(a)** Coronal slice of a representative example animal with GCaMP6f signal in green (left), FoxP2 and Calretinin staining in blue and red, respectively (middle) and the corresponding merged image (right). **(b)** Lens locations of the animals (*n* = 10) displayed in all experiments except for animals in simultaneous contrast experiments in Fig. 4e-h and Extended Data Fig. 5 shown in **(c)** (*n* = 4).

**Extended Data Fig. 3:**
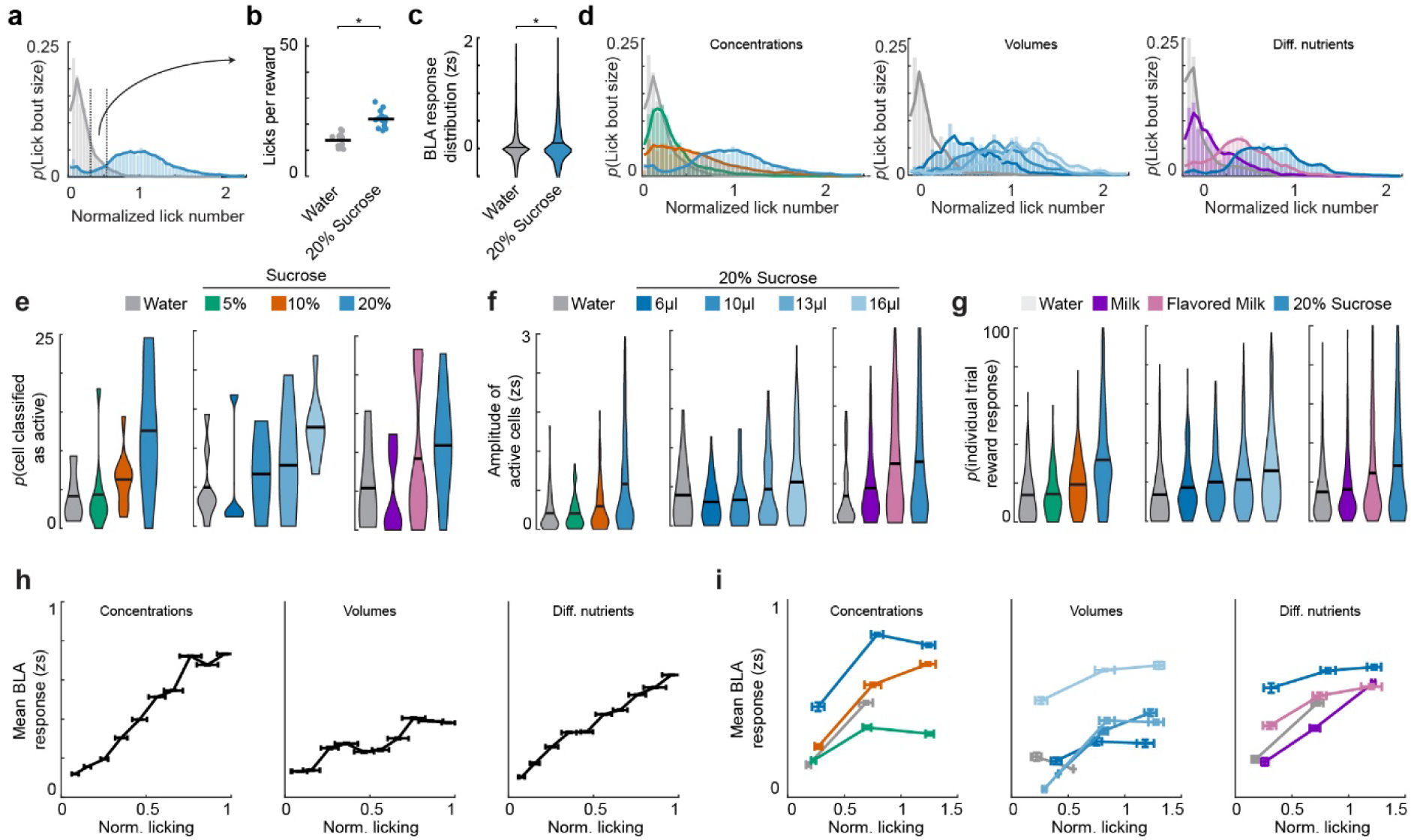
Despite stable stimulus representation across lick bout sizes, value scaling arises across different rewards through differential strength of activation of neurons. (a) Distribution of licks for the different concentrations, dotted lines indicate selected range (0.25 - 0.75 norm. licks) of trials, with corresponding quantification of mean licks per reward (*p* = 1.22*10^(−07), *n* = 10 (*animals*) - *sessions* = 4) (b) and the mean BLA response (*p* = 1.67*10^(−05), *n* = 780 (*neurons*) - *animals* = 10, *sessions* = 4) (c). (d) Distribution of lick bout sizes across the three different session types Concentration (left), Volume (middle) and Diff. nutrients (right). (e) Mean amplitude of the BLA across all stimuli split according to the number of licks during the trial across the concentration sessions (left), volume sessions (middle), and the Diff. nutrients session (right). (f) Same as (e), however with split reward responses.

**Extended Data Fig. 4:**
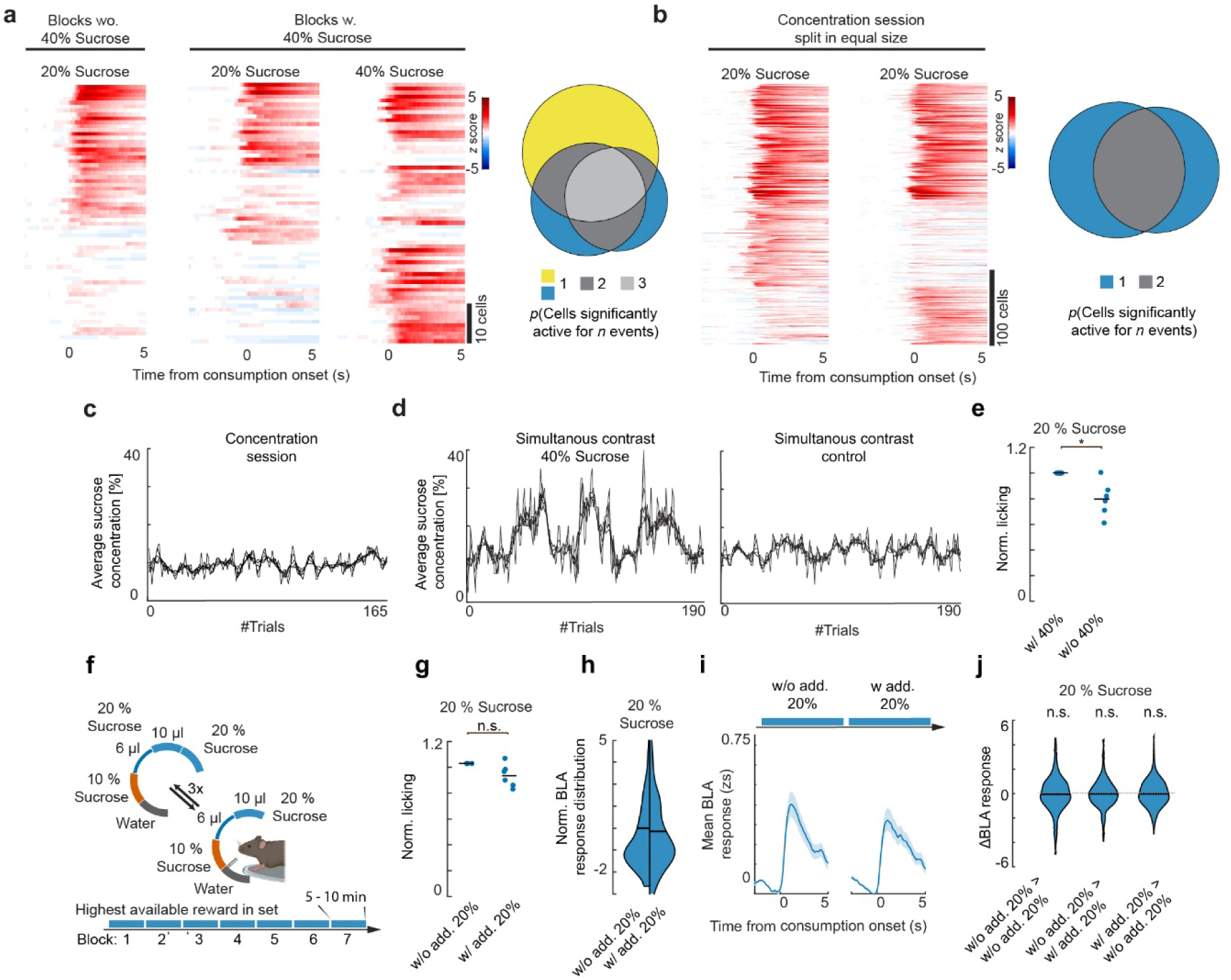
Overlap of population scaling sucrose response, reward history quantification, licking response effect, and control condition showing simultaneous contrast is only apparent for higher valued reward. (a) Neurons responding to 20% sucrose and 40% sucrose in blocks without 40% sucrose (left), with 40% (middle) and corresponding quantification of overlap (right). (b) Analog plot to (a), generated by splitting trials in the concentration sessions into two subsets and visualizing the same analysis (left) with corresponding quantification (right). (c) Average sucrose concentration plotted for the last 2, 4, 6, 8 and 10 trials in the different concentration experiment across the experiment presented in Fig. 1 and 2 (d) Average sucrose concentration plotted for the last 2, 4, 6, 8 and 10 trials in the simultaneous contrast experiment for the experimental group (left, related to Fig. 4) and control group (right, Fig. 3) across the experiment (e) Lick behavior for 20% sucrose during the different phases of the 40% sucrose session displayed in Fig. 4e-h (*p* = 0.011, *n* = 6 (*animals*) – *sessions* = 3) (f) Schema of control experiment with alternating blocks including additional 20% sucrose trials (g) Lick behavior for 20% sucrose during different blocks of the control experiment (*p* > 0.05, *n* = 6 (*animals*) - *sessions* = 3) (h) Split violin plots without (left) and with additional 20 % sucrose present in the current block (right) (*p* = 0.27, *n* = 175 (*neurons*), *animals* = 4, *sessions* = 3) (i) Time-resolved mean BLA response to 20% in blocks without (blocks 1 & 3, 5), and the blocks with (blocks 2 & 4, 6) additional 20% sucrose. (j) Quantification of the ΔBLA response between 20% blocks (blocks 1-3, 3-5, 5-7, left), transitions from 20% blocks to additional 20% blocks (1-2, 3-4, 5-6, middle) and transitions from additional 20% blocks to 20% blocks (2-3, 4-5, 6-7, right) (*p* > 0.5; *p* > 0.5; *p* > 0.5; *n* = 326/350/345 (Δ*Neuron activity*), *animals* = 4, *sessions* = 3, *blocks* = 3, *neurons* = 175). Shaded regions depict SEM.

**Extended Data Fig. 5:**
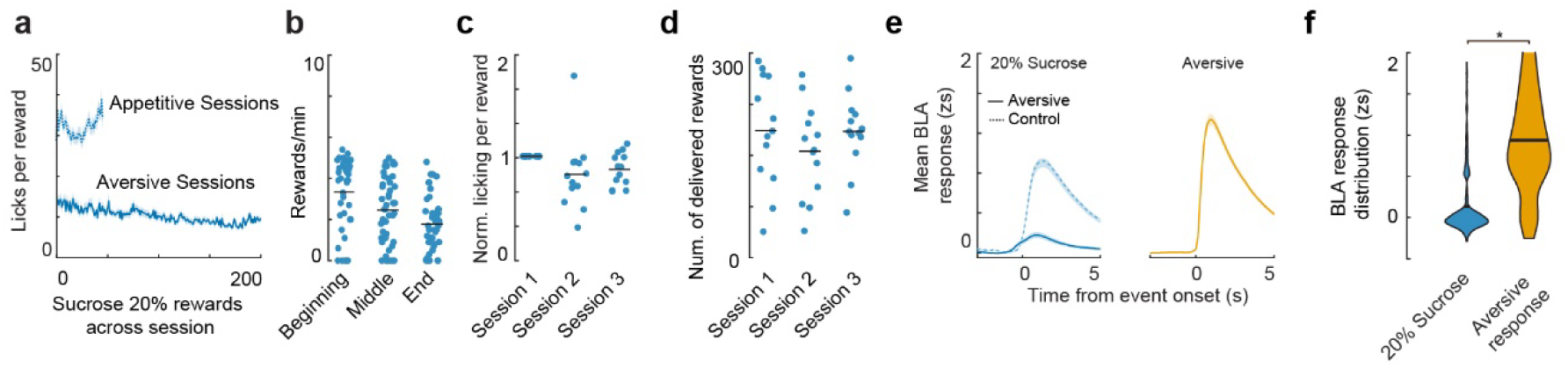
Within, but not across-session adaptation of lick bout size, motivation and aversive BLA response. **(a)** Average number of licks for 20% sucrose during appetitive and aversive sessions (*n = 10* (*animals*), appetitive *sessions* = 10, aversive *sessions* = 3) **(b)** Average number of rewards taken during the first middle and last 10 minutes of the session (*n* = 39; *animals* = 13, *sessions* = 3) **(c)** Normalized average number of consummatory licks for 20% sucrose during the different aversive sessions (*n* = 13 (*animals*)) **(d)** Number of 20% sucrose rewards delivered during the different aversive sessions (*n* = 13 (*animals*)). **(e)** Average response of appetitive responses (left) in the aversive session (solid line) and 20% sucrose during the concentration session (see Fig. 2B, dashed line), contrasted with the response of neurons to aversive events (right), with corresponding quantification (*p* = 3.59*10^-(34), *n =* 258 (*neurons*), *animals* = 10, *sessions* = 3) **(f)**. Shaded regions depict SEM.

## Methods

### Animals

All animal procedures were performed in accordance with institutional guidelines and with current European Union guidelines and were approved by the Veterinary Department of the Canton of Basel-Stadt, Switzerland. Male C57BL6/J (Envigo, in house breeding) were housed on a 12-h light/dark cycle and had *ad libitum* access to food and water in their home cage. Environmental enrichment was provided in the form of a running wheel, bedding material, a cardboard tunnel and a shelter. Animals were between 8 and 12 weeks at the time of experiments at which time they were separated for the duration of the experiment. All experiments were performed during the light cycle. No statistical methods were used to pre-determine sample size and due to the automatic scoring and extraction of behavior the experimenters were not blinded to the experimental conditions.

### Surgical procedures, viral vector injections and GRIN lens implantation

One day prior to the surgery and three days after the animals received drinking water supplemented with the nonsteroidal anti-inflammatory drug Carprofen (Rimadyl, 67 μg/ml, 10 mg/kg, Pfizer) to provide pain relief without the necessity of injections. Before anesthesia, buprenorphine (Temgesic, Indivior UK Limited; 0.1 mg/kg body weight (BW)) was injected subcutaneously 30 min before the surgery. Anesthesia of mice was induced with 5% and maintained with 1-2% isoflurane (Attane, Provet) in oxygen enriched air (Oxymat 3, Weinmann), animals heads shaved and placed in a stereotactic frame (Model 1900, Kopf Instruments). Mice were kept on a heating pad controlled by a feedback-based DC temperature control system (FHC) and received an ocular gel to prevent drying of the eye’s surface (Viscotears, Novartis). Before the first incision was made the skin above the skull was injected subcutaneously with a local anesthetic (1:1 mixture of Lidocaine: 10mg/kg, Bichsel and Ropivacaine: 3mg/kg; Naropin, AstraZeneca). A hole of 1.3 mm diameter was drilled using a surgical drill (Kopf Instruments) at the following coordinates: AP −1.6 mm (from bregma), ML −3.35/+3.35 mm (from bregma) and an adeno-associated virus (AAV2/5.CaMK2.GCaMP6f - 500 nl, University of Pennsylvania Vector Core, UPenn) unilaterally injected into the BLA using a precision micropositioner (Model 2650, Kopf Instruments) and pulled volume-calibrated glass capillaries (Drummond Scientific, Cat.-No. 2-000- 001, tip diameter about 30 μm) connected to a Picospritzer III microinjection system (Parker Hannifin Corporation) at the following depth: DV −4.2 mm (from pia). Following the retraction of the pulled glass capillary, a hollow injection needle with 1.1 mm diameter was lowered to −4.3 mm (from pia) and immediately retracted before we lowered a 1 mm gradient refractive index (GRIN) lens into the skull (Inscopix, part ID: 1050-002177) using the micropositioner and fixed at −4.2 mm (from pia) using a UV-curing glue (Henkel, Loctite 4305). The surface of the exposed skull was scratched, covered with Vetbond (3M) and a custom-built titanium head-bar fixed using dental acrylic (Paladur, Heraeus). To protect the surface of the GRIN lens before the experiment it was covered with a rapid-curing silicone elastomer (KauPo).

### Immunohistochemistry

Following the completion of behavioral paradigms, the brains of the mice were post-processed for histological analyses. Mice were anesthetized with an intraperitoneal injection of an anesthetic mixture (250 mg/kg ketamine and 2.5 mg/kg medetomidine) to achieve deep anesthesia. The mice were then perfused transcardially with 0.1 M phosphate-buffered saline (PBS, pH 7.4) for 2 min followed by 4% paraformaldehyde (PFA) in PBS for 40 min. Then, the brains were removed from the skull and cut into 80 or 100 μm coronal slices using a vibratome (VT1000S). Perfusion-fixed sections containing the amygdala were subsequently washed in 0.1 M phosphate buffer (PB, pH 7.4) and Tris-buffered saline (TBS, pH 7.4) and blocked in a solution of 1% human serum albumin (Sigma-Aldrich) and 0.1% Triton-X (Sigma-Aldrich) in TBS and then incubated in a mixture of primary antibodies containing 0.1% Triton-X for 48-72 h. This was followed by extensive washes in TBS, and incubation in the mixture of appropriate secondary antibodies overnight. Following this, sections were subsequently washed in TBS and PB, dried on slides and covered with Aquamount (PolySciences). Sections were scanned with a Zeiss Axio Scan Z1 Slide scanner equipped with a Zeiss objective (Fluor 5×/0.25) to confirm the expression of the different viruses and GRIN lens implantation sites. GRIN lens placements were matched against a mouse brain atlas (Allen Brain Atlas)^38^.

### Behavioral protocols and recordings

Five days prior to the start of the experiment animals were food restricted and maintained at 85% of their ad libitum food body weight throughout the experiment. The feeding schedule was maintained throughout the experimental process and happened immediately after the daily experimental session was completed. Acute water restriction was performed on non-food restricted animals and constituted a 24 h period of water removal before the experiment was conducted. After the experiment finished animals were again given *ad libitum* access to water.

### Head-fixed paradigms

Two to six weeks after the surgical procedure, animals were placed under a two-photon microscope (Ultima Investigator, Bruker) to check for expression of the calcium indicator GCaMP6f. If animals had sufficient expression, they were put under food restriction (s. *above*) and the experiment commenced five to seven days afterwards.

First, mice were habituated to head-fixation and reward delivery in two brief 15 min sessions. Rewards were delivered using up to five NE-1000 syringe pumps (NewEra Pumpsystems Inc, Milan) controlled by an analog signal output from the two-photon microscope (Ultima Investigator, Bruker). The output pattern was generated using a custom Matlab 2017 (Mathworks) script that randomized deliveries and inter-trial intervals (ITI’s) and generated individual .txt files used by the system to control the voltage supplied to pumps that control the deliveries through a RS232 to BNC cable. Following habituation, mice underwent several sessions of each type (up to 4, separated by 24h, Fig. 1, 2, 3, 4 a-f, 5) in which the rewards were changed (order: concentrations, volumes, different nutrients, aversive, water restriction). All sucrose rewards were mixed according to the specified concentration of sucrose (Sigma-Aldrich) and delivered with different volumes. Milk (whole-milk, COOP Switzerland) and flavored milk (Emmy Energy Milk Vanilla, COOP Switzerland). Experiments pertaining to Fig 4e-f were performed on an additional set of mice that followed a slightly altered protocol (s. *below*). In each session rewards were delivered through a single custom-built spout that had five independent openings, one connected to each pump. Spout licks were detected when the mouse closed a circuit between its metal head-bar and the spout (P. Buchmann, P. Argast, J. Hinz - custom built). Conductance from the headbar to the mouse was facilitated by an ultrasound gel bridge to the neck skin. Additionally we monitored the running speed of the animals on the saucer wheel using a rotary encoder (Yumo rotary encoder E6B2- CWZ3E: 1024 pulses per rotation) and the pupil dilation with a single monochrome camera at 60 Hz (The Imaging Source, part ID: DMK 22BUC03) through ICcapture (The Imaging Source) or lick trajectories and pupil dilation with multiple monochrome cameras (FLIR, part ID: BFS-U3-16S2M-CS) at 194.1 Hz through Bonsai (version 2.3.1)^39^ and Python (version 3.8.8, based on https://t.ly/FHXfO) for Fig. 1f-h. To allow for a good dynamic range in pupil size, we installed a custom build 415 nm UV back light (ILS ILH-XC01-S410-SC211WIR200 filtered through a 445/45nm bandpass) and adjusted the light power to result in a medium sized pupil at the beginning of each experimental session.

During the appetitive open-loop sessions, rewards were delivered at random intervals (ITI 15 - 25 s, equal probability) and no structure imposed on the distribution of rewards. During experiments pertaining to Fig. 4e-h and Extended Data Fig. 4, rewards were organized in blocks of 25/35 (either including or excluding additional reward), where each block contained the same number of rewards, randomized within the block, and delivered at random ITIs (15 - 25 s, equal probability).

During the aversive sessions, reward delivery was contingent on the animals licking and therefore licking was additionally processed by a RP2.1 real-time processor (Tucker Davis). Rewards were delivered when the animal had licked 3 times and didn’t incur a reward time-out (8 s after the last reward was delivered).

### Behavioral Analysis

#### Data extraction

Behavioral data was synchronized with the two-photon imaging frames using the two-photon system’s GPIO channel voltage recording (Ultima Investigator, Bruker) at 1 kHz. The running speed was determined from the rotary encoder signal by normalizing to the running saucer wheel circumference at the mouse position. Video camera frames were timestamped by recording the frame trigger signal. Pupil dilation, whisking power, and lick trajectories were extracted from the video camera recordings via DeepLabCut (DLC)^40^, using a network trained on manually labeled sample frames (8 points around the pupil, 4 points around the whisker pad, 3 points on the tongue, 3 points on the lick spout).

#### Data Analysis

To ensure that we only included actual licks and not spout contacts by other body parts, like hand digits, were detected, we used the length of contact to exclude other events. Tongue contacts were the most common spout interaction and had a stereotypic duration. We therefore fitted a normal distribution to the duration of all detected spout contacts and excluded events that were longer or shorter than 3*std of the mean contact duration. Given the delivery speed of the syringe pumps, we defined the first rewarded lick to be the first lick 100 ms after pump onset. This definition comes with some level of uncertainty so that, given the average lick frequency of 6.8 Hz, the first rewarded lick can in some cases be 150 ms earlier or later than the one we defined. To calculate the lick bout size, we counted the number of consecutive licks that were not interrupted for more than 500 ms. We normalized these lick bout sizes to the average within animal, 10 µl, 20% sucrose lick bout size. To exclude an impact of reward prediction error on the observed neuronal signals we checked whether pupil dilation was impacted by positive or negative reward difference. To this end we fit the distribution of pupil dilations and checked whether there was a significant deviation from the mean of the corresponding distributions for positive and negative reward shifts (*see* Extended Data Fig. 1c-e). To investigate the motor patterns during consumption we analyzed running speed and pupil dilation (*as described above*), as well as whisking power and lick trajectories. Whisking power was defined as the average pixel value change within the whisker pad area. To compensate for potential lick speed differences, lick trajectories were interpolated to the same number of coordinates per lick using a modified Akima cubic Hermite interpolation (Matlab, interp1, option makima, upscale factor 10) from the DLC tracked tongue tip trajectory around the electrically detected licks times. Lick trajectory similarity was defined as the mean Procrustes distance (Matlab, procrustes, no reflection) between all compared lick trajectories.

### Two-photon Imaging

Imaging was conducted using a two-photon microscope (Ultima Investigator, Bruker) during all behavioral sessions on three planes separated by 70 - 80 μm using a 400 μm mechanical piezo (Bruker) with bi-directional scanning at a volume rate of 8.5 Hz and a spatial resolution of 256 by 256 pixels through a water immersion objective (16x, 0.8 numerical aperture (NA), Nikon). Ultrasound gel (G008, FIAB spA) was used to interface the objective and the GRIN lens. Excitation light was provided with a mode-locked laser system operating at 920 nm, 80-MHz pulse rate, 120 fs pulse width (Insight X3, Spectra Physics). We used standard Bruker PMT (photomultipliers, Hamamatsu Photonics) for light detection. For the same mice, imaging parameters were kept the same across repeated behavioral sessions and recorded using Prairie View (versions 5.2 - 5.6, Bruker).

#### Data extraction

We extracted the images using custom software written in Matlab 2017 (Mathworks Inc.) available on GitHub^28^. Briefly, after loading individual *.tif* and converting them to .mat files they were mapped to RAM and rigid motion correction performed using the Matlab implementation of *NoRMCorre*^41^. Animals that displayed non-rigid motion were excluded in a manual intervention step and ROIs identified using the Matlab implementation of *CNMF*^42^. The resulting ROIs were initially manually sorted and these manually sorted components used to train an SVM classifier that was subsequently used to re-sort all previous components and all future datasets. After the automatic sorting procedure, we excluded components that had high temporal correlation within and across planes when they were in close spatial proximity. We then aligned the remaining components across sessions by first performing rigid field-of-view alignment using landmark registration and then aligning the components using the cross-day alignment code of the package *CIAtah*^43^. Next, we combined the data for the multiplane acquisition that needed separate analysis and took the timestamp of the middle image as the timestamp for the three planes (± 4 ms).

#### Data analysis

All analysis of calcium data was performed on deconvolved traces (variable *C_dec*) extracted through *CNMF* and performed with custom code written in Matlab 2017b (Mathworks, code available on Github).

The first step was to align the first rewarded lick, we obtained by aligning the licking to the reward delivery (see *behavioral analysis* for details), to the imaging. Therefore, we used the timestamp of acquisition for each image and found the closest image recorded to the first rewarded lick and extracted a window of −3 s to +5 s around this timestamp for further processing. As pointed out previously there are some uncertainties about the first rewarded lick and given the volume imaging rate of the two-photon system of 8.5 Hz and the associated confounds we estimate that the actual onset of neuronal activity associated with the reward consumption is in the range of a max./min. of 250 ms of what we state as the time of consumption. This means that there is a jitter around zero of about ± 2 frames most likely explaining imperfect alignment of the activity to time point 0 s.

To present peri-stimulus time histograms (PSTH) across the paper we computed local z-scores by normalizing each individual trial. To this end we subtracted the mean activity of each baseline before consummatory onset from the whole extracted period and divided it by the variance of the whole neuronal trace. Through this process we avoided the issue of division by zero, which often happens when using deconvolved traces and corrected for the overall activity level of the neuron. The mean activity for neurons displayed in the paper is the mean of the normalized individual presentations. To calculate the *p*(Ca2+-transients) we used the estimated spike variable, binarized it and calculated the mean and SEM around the time of consumption of the different rewards for each neuron. Displayed is the mean across neurons and the threshold derived by calculating the mean and 3 standard deviations of the baseline preceding the consumption onset. To quantify matched lick bout trials, we identified sessions that had at least 3 repetitions for each of the stimuli within the normalized lick range of 0.25 - 0.65 of the mean 20% sucrose response. These trials were then pooled across animals and used for quantification and visual display. We calculated the PV-correlation by extracting the mean response (s. above for definition), for individual trials across all recorded neurons resulting in a *1d* vector, which we used to calculate the Pearson correlation between each single reward presentation. We subsequently sorted these correlations by reward identity. To calculate the correlation of single neurons with all stimuli or the individual stimulus regressors, we correlated the average activity of significantly active neurons during each trial with either a regressor containing all ones or ones for one stimulus and zeros for the rest. To quantify the mean activity for specific lick bouts, we calculated the mean response of active neurons in the specific lick bout, either split according to stimulus identity (Extended Data Fig. 3i) or summed over all stimuli (Extended Data Fig. 3h). During simultaneous contrast experiments we calculated the change of the 10 µl 20% sucrose response between the different blocks by subtracting the mean response of neurons with at least 3 measured events and calculating whether these differences were deviating from 0. Concurrently we also calculated the relative BLA response in Fig. 5g (between different rewards) and Fig. 4d, where we subtracted the mean BLA response from the indicated rewards (e.g., 10 µl Sucrose response during Session II - mean BLA population response of 10 µl Sucrose during Session I).

### Antibodies

We used DAPI (Thermo Fisher Scientific, Cat# D1306), Goat-anti-Calretinin (Dilution 1:5000, Swant, CG1), Chicken-anti-GFP (Dilution 1:2000, Lifetech, A10262), Rabbit-anti-vAchT (Dilution 1:3000, Synaptic Systems, 139103) and Mouse-anti-FoxP2 (Dilution 1:3000, Millipore, MABE415) as primary Antibodies in the identification of lens positions. We visualized them with the following secondary antibodies: Donkey-anti-Chicken, Cy2 conjugated (Dilution 1:1000, Jackson Immunoresearch Inc., 703- 225-155), Donkey-anti-Mouse, 488 conjugated (Dilution 1:500, Molecular Probes, A21202), Goat-anti-Chicken, 488 conjugated (Dilution 1:1000, Molecular Probes, A11039), Donkey-anti-Rabbit, Cy3 conjugated (Dilution 1:1000, Sigma, AP182C), Donkey-anti-Goat, 568 conjugated (Dilution 1:500, Molecular Probes, A11057) and Donkey-anti-Mouse, 647 conjugated (Dilution 1:500 – 1:1000, Molecular Probes, A31571). The specificities of the primary antibodies were extensively tested, using knock-out mice if possible. Secondary antibodies were extensively tested for possible cross-reactivity with the other antibodies used, and possible tissue labeling without primary antibodies was also tested to exclude auto-fluorescence or specific background labeling. No specific-like staining was observed under these control conditions.

### Statistical analyses and data presentation

Statistical analysis was performed with Matlab 2017b and Matlab 2022a (Mathworks). No methods were used to predetermine sample size. The sample sizes used here match those usual for the field. Experimenters were not blind to conditions, but all sorting and data extraction performed automatically, and sorting algorithms found on GitHub. We excluded animals if at least one of the following conditions were met: **(i)** GRIN lens placement outside of the BLA, (ii) non-rigid motion artifacts in the recordings, (iii) insufficient GCaMP6f expression or (iiii) insufficient interaction with the spout after habituation. All data are expressed as mean ± SEM unless stated otherwise. All data is always included in the visualization, no data was excluded from the analysis. Datasets were acquired from mice stemming from multiple litters and were reproducible. The statistical significance threshold was set at *p* < 0.05 and significance indicated by *\** and the exact *p*-value reported in the text and/or figure legend. For figure display, traces were normalized to represent the local z-score as described under the *Data analysis*. We display multiple comparison corrected *p*-values using the Bonferroni correction. We used repeated measures one-way analysis of variance (ANOVA) for more than two groups and a Tukey-Kramer test if individual differences needed to be compared. When less than 3 groups were compared, paired or unpaired student’s *t*-tests were used. A Fisher’s exact test was conducted to compare proportions between two groups.

